# Spatiotemporal dynamics of synthetic microbial consortia in microfluidic devices

**DOI:** 10.1101/590505

**Authors:** Razan N. Alnahhas, James J. Winkle, Andrew J. Hirning, Bhargav Karamched, William Ott, Krešimir Josić, Matthew R. Bennett

## Abstract

Synthetic microbial consortia consist of two or more engineered strains that grow together and share the same resources. When intercellular signaling pathways are included in the engineered strains, close proximity of the microbes can generate complex dynamic behaviors that are difficult to obtain using a single strain. However, when a consortium is not cultured in a well-mixed environment the constituent strains passively compete for space as they grow and divide, complicating cell-cell signaling. Here, we explore the temporal dynamics of the spatial distribution of consortia co-cultured in microfluidic devices. To do this, we grew two different strains of *Escherichia coli* in microfluidic devices with cell-trapping regions (traps) of several different designs. We found that the size and shape of the traps are critical determinants of spatiotemporal dynamics. In small traps, cells can easily signal one another but the relative proportion of each strain within the trap can fluctuate wildly. In large traps, the relative ratio of strains is stabilized, but intercellular signaling can be hindered by distances between cells. This presents a trade-off between the trap size and the effectiveness of intercellular signaling, which can be mitigated by controlling the initial seeding of cells in the large trap. These results show how synthetic microbial consortia behave in microfluidic traps and provide a method to help remedy the spatial heterogeneity inherent to different trap geometries.

## Introduction

Synthetic gene circuits in bacteria have traditionally been constructed and characterized in single strains.^1–3^ In nature, however, microorganisms are commonly found in consortia of multiple interacting strains and species. This allows for population-level phenotypes that improve survival in environments that fluctuate across time and space.^4–6^ Synthetic biologists have begun designing gene circuits distributed across two or more communicating microorganisms to create synthetic microbial consortia that exhibit phenomena difficult to engineer into single strains.^7–10^ Synthetic microbial consortia allow complex pathways to be split among specialized strains, reducing the metabolic burden on each cell.^11^ Optimizing each step of the divided pathway in separate strains is simpler than optimizing an entire pathway within a single cell.^12^ In addition, synthetic consortia allow for the study of population-level phenotypes dependent upon intercellular and inter-strain communication.^4^

Microfluidic devices have long been used for assessing gene expression and growth of engineered bacteria as they allow for continuous growth and visualization over time.^13–16^ However, complications can arise when multiple strains are grown together within a microfluidic device. For instance, spatiotemporal strain ratio fluctuations can affect the relative amounts of intercellular signaling molecules and complicate the analysis of intercellular interactions.^10,17^ Another complication arises in microfluidic devices with large cell-trapping regions: Initial seeding and subsequent growth patterns can lead to large areas containing just a single strain. Because intercellular signaling molecules have a limited diffusion range in the trap and can be lost to open boundaries, cells in one portion of the trap may not communicate directly with cells in another. Therefore, if the strains within a consortium are not well mixed, signaling between the strains will occur only near their interfaces. However, in small traps, even well-mixed populations are subject to severe strain ratio instabilities, which can lead to the complete loss of one or more strains within a consortium.

In this study, we compared two common microfluidic devices (Figure 1) by characterizing spatiotemporal population and signaling dynamics in consortia of two nearly identical engineered strains of *E. coli.* We found that temporal fluctuations of strain ratio in small trap experiments could become severe, whereas large traps stabilized the ratios. However, increased spatial separation of strains in large traps could limit intercellular signaling. We demonstrate how to mitigate this problem by reliably tuning the spatial mixing of the two strains to ensure that cells of each type are close enough to exchange signaling molecules. Our results demonstrate the value of microfluidic devices as a tool for studying the dynamics of synthetic microbial consortia, despite the challenges they present.

**Figure 1:**
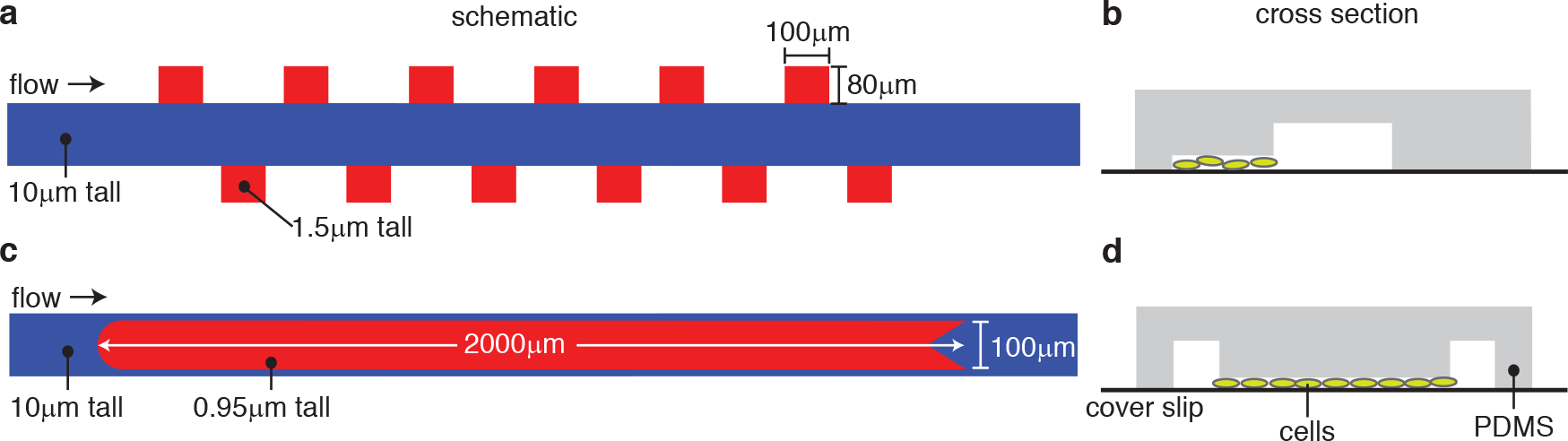
Designs of the microfluidic devices. **(a)** Schematic of the “hallway” device. The blue region is the tall flow channel and the red regions are the shorter cell-trapping regions. **(b)** Cross section of the hallway trap device shown with cells in the trapping region. **(cd)** Same as (ab), but for the “open” device. See text for more details.

## Results and Discussion

We examined synthetic microbial consortia in microfluidic devices with two distinct types of cell traps, as shown in Figure 1 (also see Figure S5). The “hallway” device^10^ contains several traps (80 μm × 100 μm) on either side of a media flow channel (Figure 1a,b). Variations of the hallway device have been used in several studies to increase experimental replicates or to allow for multiple communicating populations.^10,18,19^ Because the traps in this device are small compared to the mean diffusion distance of HSL signaling molecules in one cell growth period,^20^ and because there are three closed walls (reflecting boundaries), cell-cell signaling molecules can quickly diffuse throughout the chamber with little to no spatial gradients (Figure S4).

However, due to small effective trap area, each hallway trap can only contain a small population,^10^ which, as we will see, can amplify the severity of temporal strain ratio fluctuations. In contrast, the “open” device (no closed walls) has a single, longer trap (100 μm × 2000 μm) that is centered within the main flow channel (Figure 1c,d). Similar trap configurations have been used to assess the spatiotemporal dynamics of large populations.^18^ Due to its larger trapping area, this trap can sustain a much larger population than an individual trap in the hallway device. However, the spatial degradation of cell-cell signaling becomes important: Diffusing molecules have a limited range compared to the spatial extent of the trap (see Figure 6). Note that the two types of traps have different trap heights to allow for proper cell seeding in each design (see Methods).^18^

### Non-communicating strains

We first studied how the population strain ratio is affected by the size and geometry of the two trap designs. When referencing size we are referring to the area of the cell-trapping region. For example, the open trap has a larger cell-trapping area than the hallway trap and can hold approximately 70,000 cells, compared to approximately 3,000 in each hallway trap. When discussing geometry we are referring to the different properties of the trap, such as open versus closed walls, trap height, and location in relation to the media flow channel.

For example, the open trap geometry consists of a trapping region located within the media flow channel, a trap height of 0.95 μm, and no closed walls. In contrast, the hallway trap is located off to the side of the flow channel, is 1.5 μm in height, and has three closed walls.

To compare population growth in these two traps, we used two non-communicating strains of *E. coli*, each expressing a different fluorescent protein for identification; we refer to these as the “control” strains. These strains are the same BV25113 derivative transformed with a plasmid that constitutively expresses the gene encoding either yellow or cyan fluorescent protein (*sfYFP* or *sfCFP*). Because *sfYFP* and *sfCFP* differ by only a few point mutations,^21,22^ the growth rate, size, shape, and biofilm production of the two strains were identical (Figure S6), and neither had a fitness advantage over the other in the microfluidic device. We mixed cultures of both strains and seeded many cells into the traps to increase the probability of simultaneously capturing both strains. We then mounted the microfluidic device onto an inverted microscope and monitored phase contrast, yellow fluorescence and cyan fluorescence channels every 6 minutes for up to 20 hours. To quantify the ratio of the strains as a function of time, we measured the total number of intracellular pixels containing either yellow or cyan fluorescence within a boolean mask computed for each frame for each image acquisition frame over time (see Methods).

### Hallway traps

As shown in Figure 2a,b, strain ratios exhibited large temporal fluctuations when grown in the hallway trap. *Boyer et al.*^23^ demonstrated that a growing colony of bacterial cells in a three-walled trap is prone to a *buckling instability* when cells at the back of the trap become aligned with the back wall (due to the finite width of the back wall and the continuous axial growth of the rod-shaped bacterial cells, the cells buckle from the large axial stress that results). In our two-strain experiments, the buckling instability can translate directly to a strain ratio instability because cells at or near the closed back trap wall serve as progenitor cells for the entire trap population: Cells continuously grow and expand while pushing each other generally in random directions. However, only the open end of the trap is available for cells to exit the trap. Thus, the local strain ratio near the back of the trap effectively determines the strain ratio in the entire trap.

**Figure 2:**
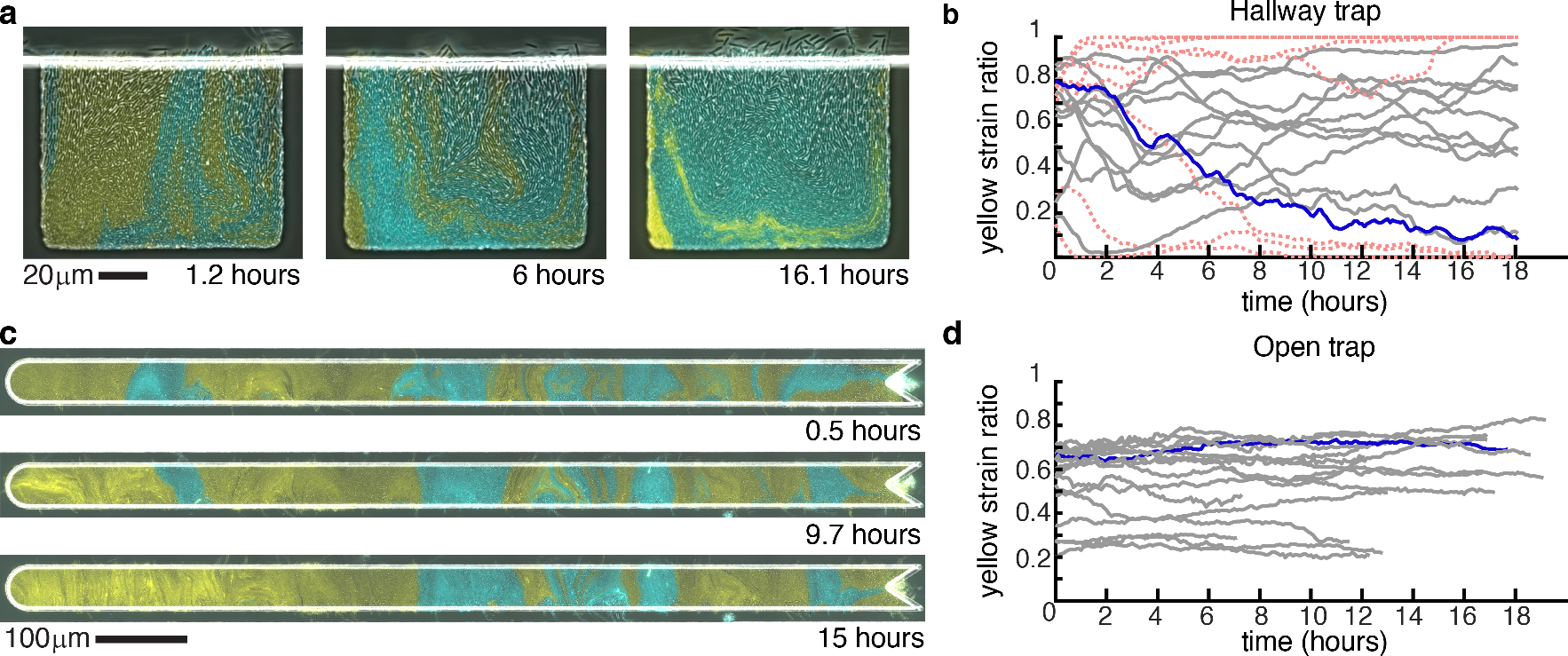
Strain ratio time series. **(a)** Images of the two control strains grown in a hallway trap over time. **(b)** The ratio of the yellow strain in all experiments performed in the hallway trap with the experiment from (a) in blue. Red dotted lines signify traps that lose one strain completely. **(c)** Images of the two control strains grown in the open trap over time. **(d)** The ratio of the yellow strain in all experiments performed in the open trap with the experiment from (c) in blue. Time 0 was defined as the first image at which the entire trap was filled with cells.

Assuming that the two strains in our consortium on average push each other out at equal rates via volume exclusion, we conclude that strain ratio fluctuations can result from the stochastic effects of buckling instabilities influencing the back-area ("mother" cell) positions in the hallway trap. These fluctuations alter the composition of the population over time and, importantly, can impact the intercellular signaling dynamics of synthetic microbial consortia by impacting the resulting balance of signaling molecules.

### Open-walled traps

Reducing strain ratio fluctuation is important for maintaining communication in consortia, and we found that altering the size and boundary geometry of the trap can play a role in population stability. We grew the same control consortium in the open trap and observed more stable strain ratio trajectories than in the hallway trap (Figure 2c,d,). We found that the range of fluctuations of the yellow strain ratio differed significantly between open and hallway trap experiments (Figure 3a), (*p* < 0.01). The increased strain stability in the open device may be attributed to two main factors: the different geometries of the traps (affecting cell buckling) and an overall increase in population size (a large number effect).

In contrast to the hallway device, the open device has no closed walls and allows cells to grow in every direction, resulting in decreased axial growth pressure^24^ and an emergent alignment of the cells that is sustained in the bulk of the colony.^24–26^ When we consider the aspect ratio of the open device (see Figure 1), cells will tend to form vertical columns, with the "mother" cells at or near the middle of a column establishing the strain column identity.^27^ The lateral movement of "mother" cells in the trap (i.e. those near the vertical center) has the highest impact on strain ratio: If a cell of one strain laterally displaces a cell of a second strain, the invading cell will become a "mother" cell and thereby change the population balance by eventually occupying the entire column. Such behavior is stochastic and may include lateral flux of varied intensity in both directions.

Since the width of the open trapping region is 20:1 in ratio to its height, we expect the global strain ratio in this device to be less sensitive to random, lateral cell fluctuations when compared to the hallway device. Under an assumption that both strains are subject to similar stochastic lateral motion, the local fluctuations would be averaged over a larger area in a longer device. This is a clear advantage of the open device over the hallway device with respect to strain ratio stabilization. However, given that the growth pressure is also significantly reduced in the open device as compared to the hallway device (at the respective "mother" cell positions),^24^ we would also expect smaller strain ratio fluctuations in the open device due to relaxation of the buckling instability: Mitigation of the buckling instability directly translates to increased strain ratio stability since the "mother" cell positions are stabilized, resulting in less lateral motion at the critical strain-strain boundaries where strain ratio fluctuations can emerge.

**Figure 3:**
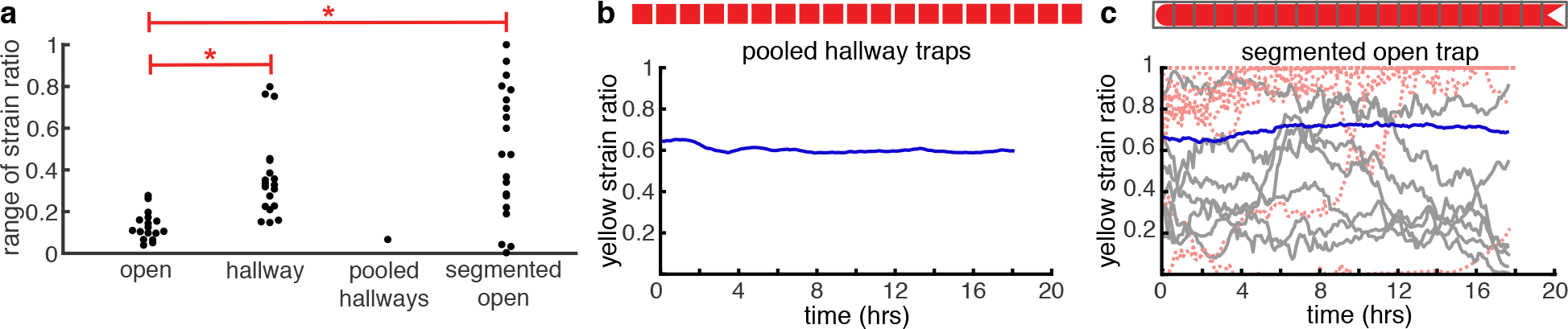
Strain ratio comparisons. **(a)** The range of the yellow strain ratio fluctuation over time is significantly different in hallway traps (Figure 2b) than in open traps (Figure 2d, *p* < 0.01, student’s *t*-test). **(b)** When pooling data from all 20 hallway traps to mimic the population size in an open trap, the pooled yellow strain ratio is more stable. The range of this data is shown in panel (a) as “pooled hallways”. **(c)** Segmenting the data from the open trap experiment shown in Figure 2c into hallway-sized segments and plotting the yellow strain ratio within each segment over time results in higher variability, compared to the overall ratio (heavy blue line). Red dotted lines are lines that go to 0 or 1. The range of yellow strain fluctuations within each segment are shown in panel (a) as “segmented open.” These fluctuations do not differ from those in hallway traps (*p* > 0.01), but do differ from fluctuations in the entire open trap (*p* < 0.01).

### Isolating trap size and geometry

Open traps can contain roughly 70,000 cells as compared to the roughly 3,000 cells in each hallway trap. We wanted to isolate the affect of population size from the affect of trap geometry on strain stability in order to better understand strain ratio fluctuations in the two devices. We began by pooling the data from all hallway traps to reach a population size close to that of the open trap and we then computed the strain ratio as a function of time.

The pooled hallway data resembles that of the open trap in population stability (Fig. 3a,b). We also segmented the open trap data from Fig. 2c into sections of the same width as a single hallway trap. The strain ratio within each segment was less stable than that of the overall open trap. We obtained similar results when we segmented other open trap experiments (Fig. S7). The range of fluctuations of the yellow strain in the segmented open trap were no different than in the hallway traps (*p* > 0.01, Figure 3a,c), but differed from range fluctuations in the entire open trap (*p* < 0.01). These comparisons suggested that population size was a key factor in analyzing strain ratio fluctuations in these devices. However, this was not suffi cient to characterize the role of trap size versus the role of trap geometry with respect to strain ratio stability. To gain an increased understanding of the spatiotemporal dynamics of strain ratio and the mechanisms of its stability in these types of microfluidic devices, we manufactured additional versions of each trap with varying trap lengths.

To compare different population sizes for each device type, we fabricated open traps of four additional lengths: 100, 225, 500, and 1000 μm, and one additional hallway trap, 2 mm in length (Fig. 4a). Using these additional trap sizes, we could also directly compare each original device with that of the other type, but with comparable cell-trapping areas and number of cells. We grew the same control strains in these additional devices and computed the range of the strain ratio time series over the entire experiment (each experiment required ≈ 18 hours, see Fig. S8). We computed the range by subtracting the maximum and minimum of the strain ratio time series in each experiment (we found this measure to be a good indicator of variability of the strain ratio across experiments and devices; we also computed the variance of the strain ratio time series and obtained similar results, see Fig. S9).

In Figure 4b we show the mean (averaged over 6-8 experiments each) of the strain ratio range versus the area of an individual cell-trapping region in each device.

The data for the open traps show that the mean strain ratio range rapidly decreases with increasing trap size, which is expected if local strain ratio fluctuations are averaged out over a larger trapping area. A similar trend is indicated for the hallway traps, but the large variability in those measurements suggest that trap geometry plays a significant role in strain stability in addition to the role of population size.

We noted that the strain ratio range in the largest open trap (2 mm) was not significantly different than the that of the next smallest trap (1 mm). To determine if this was a limiting effect of increasing trap size, we examined the data more closely. We noticed that whenever a larger number of cells were seeded into a trap the strain ratio range was slightly higher than if fewer cells were seeded. Further, we generally observed that whenever more cells were seeded the number of distinct “bands” once the trap was filled was greater (here a band is a region of the trap composed of mostly one strain). An increase in strain bands can increase strain ratio fluctuations by increasing the spatial density of random lateral invasions of "mother" cell positions by an adjacent band, as explained above. Although the mechanisms of the fluctuations are likely a combination of several stochastic factors, we wanted to see if any relationship existed between the number of strain bands (which increase with increased number of seed cells) and the strain ratio variability. In Fig. S10 we reduced the data from the 2 mm open device to contain the same number of experiments (6) as for the smaller devices and also to contain a comparable number of bands (see the following section and Fig. S11). The reduced data show a reduced measure of strain ratio instability, which suggests a causal connection.

**Figure 4:**
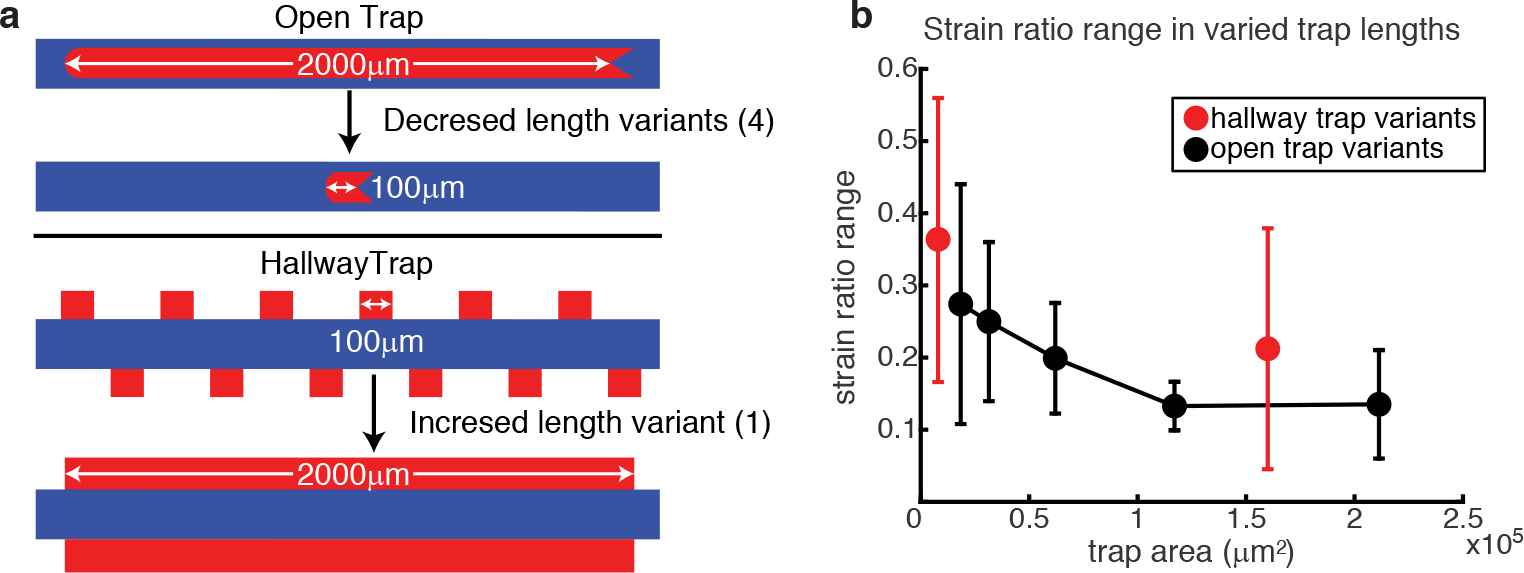
Strain ratio stability for traps of varying size. **(a)** Open traps with decreasing lengths and a hallway trap with increased length were manufactured to determine how population strain ratio dynamics depend on the size of the population in each trap. **(b)** Filled circles show the empirical mean of the strain ratio time series; ranges (*max* − *min*) for each trap size investigated; black circles: open-walled traps, red circles: hallway traps. The smallest area hallway trap and largest area open trap data points use the same data as in Fig. 2a,c. The other data points are averages of 6-8 experiments (Fig. S8). Starting with the smallest trap size, both the mean of the strain ratio ranges and their standard deviations (error bars) generally decrease with increasing trap area for open traps. For the two sizes of hallway traps, the strain ratio range mean decreases with the increased trap size, but the error bars indicate sustained, large variation across experiments in both sizes. Error bars represent standard deviation for *N* ≥ 6.

Our data support a conclusion that a larger, open trap geometry favors increased strain ratio stability in microfluidic trap experiments consisting of two (or more) strains. By increasing the cell trapping area, our experiments demonstrate a trend towards increasing stability (as measured both by range and by temporal variance) of the strain ratio time series. However, increasing the number of cells in the trap did not increase stability in all cases since the number of bands evidently can also influence strain ratio variability. A systematic study of initial cell seeding and resulting strain ratio variability is challenging since there exists little control over spatial placement of seed cells. In the next section, however, we study the effects of seed cell count on the spatial separation of strains in a two strain consortium and further explore the differences and challenges of the two microfluidic trap devices in our experiments.

### Spatial patterning and communication

Population composition needs to be stable to coherently sustain communication and distributed functionality across microbial consortia.^12^ Additionally, spatial patterns and the physical separation of strains in a multi-strain experiment are important because intercellular signaling distance can be limited. For example, while the 2mm long open trap in our experiments described above increased strain ratio stability, it also produced spatial strain distributions that could hinder inter-strain signaling due to large separations of strain types. This is in contrast to the original hallway device where signaling molecules from both strains appear to be homogeneously distributed in the trap.^10^

In our initial experiments with the longest (2 mm) open trap, the number of seed cells loaded directly impacted spatial heterogeneity of the final population. When fewer cells were initially seeded, larger bands of single strains formed in the trap (Fig. 5a). The average number of bands in a trap over time positively correlated with the number of cells initially seeded (Fig. 5b).

To better understand the relationship between the number of initial cells seeded in the trap and the number of resulting bands in the full trap, we used a one-dimensional mathematical model to represent the initial seeding of the trap as a sequence of *N* flips of a fair coin, where *N* corresponds to the number of seed cells and *H* (heads) and *T* (tails) represent the two strains (see Supplementary Information). Single-strain bands in the trap correspond to runs of heads or tails, which have a geometric distribution in this model.

Motivated by the large (20:1) aspect ratio of the 2 mm open device, we conjectured that this simple, one-dimensional model would capture the essential relationship between the number of seed cells and the resulting number of bands (see Supplementary Information for the band *number* distribution in the model), while remaining analytically and computationally tractable. Figure 5b illustrates the mean number of bands over time for each of 18 experiments. The shaded region depicts the model prediction (mean ± 3 s.d. - see Supplementary Information). As conjectured, the model is consistent with most of the experimental data (the trap with the largest number of seeded cells seems to drop off from the expected number of bands; however, we observed that in this experiment the initially seeded cells clumped together in the trap reducing the effective number of “colonies” that can form individual bands as the cells grow, see Figure S12). This consistency supports the intuitive conclusion that band formation depends on the number of initial cells seeded into the trap.

**Figure 5:**
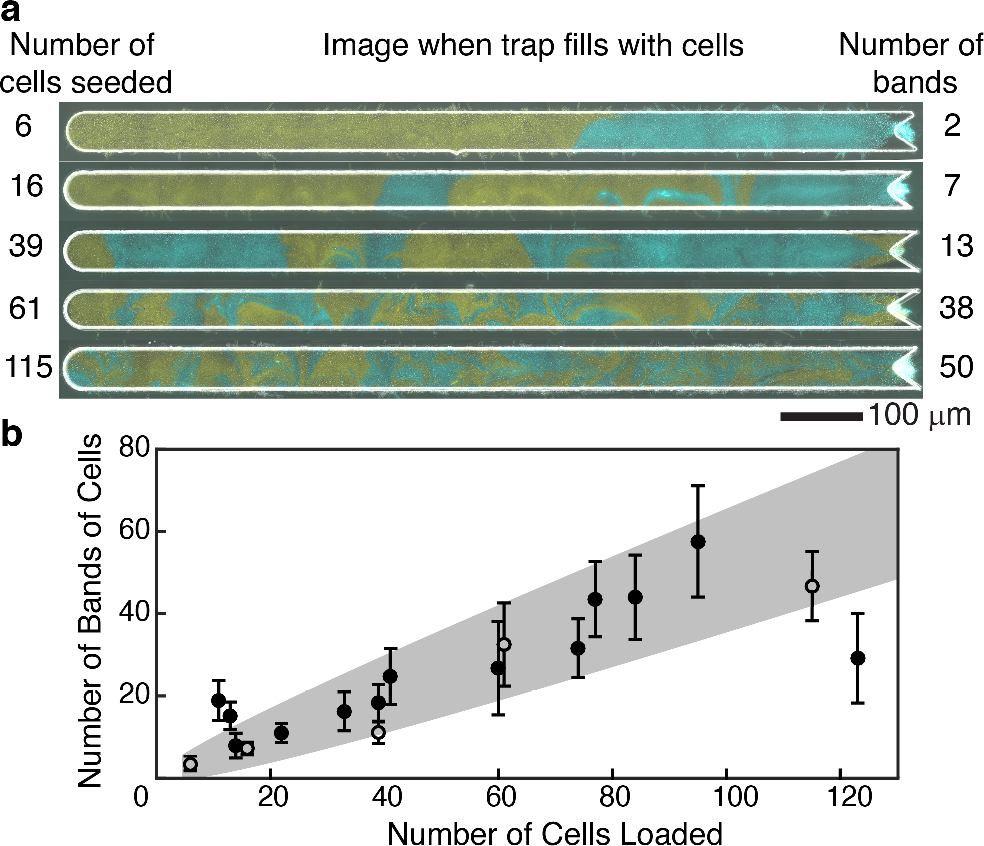
Emergent single-strain banding vs. number of cells seeded into trap. **(a)** Images from five representative experiments with two strains grown in the largest open trap at the time the trap fills completely with cells. The number of cells seeded into each trap is given on the left, and the number of bands formed when the trap fills is given on the right. The greater the number of cells seeded into the trap, the more heterogeneous the population. **(b)** The number of bands (single-strain regions) increases approximately linearly with the number of cells seeded. The points are from the same experiments shown in Figure 2. Open dots correspond to the 5 examples shown in panel (a). The predicted range of bands (mean ± 3 s.d.) as a function of seeded cells is shaded in grey (see Supplementary Information).

### Sender-receiver signaling

To assess the effect of spatial population heterogeneity on communication in microbial consortia, we grew a “sender” and “receiver” consortium in the original (2 mm) open trap (Fig. 6a). The sender strain contained a plasmid coding for constitutive expression of *sfYFP* for strain identification and constitutive expression of the quorum sensing (QS) gene *rhlI*. The RhlI enzyme produces the QS molecule N-butyryl-L-homoserine lactone (C4-HSL).^28^ This QS molecule diffuses out of the sender cells and into surrounding cells. The receiver strain contained a plasmid coding for constitutive expression of *mCherry* for identification and the C4-HSL regulated expression of *sfCFP*. Thus, varying C4-HSL capture by the receiver strain is experimentally quantified by varying measure of its CFP fluorescence.

When sender and receiver strains were co-cultured in the open trap, we observed regions of cyan fluorescence in receiver cells adjacent to bands of sender cells (Fig. 6b,c), but the signaling was of limited extent (this behavior is consistent with predictions of our mathematical model for diffusion-based signaling in this device where the concentration profile of the QS molecule along this domain is characterized by exponential decay from an idealized sender-cell boundary, see Supplementary Information).

The effective reach of QS molecules depends on the heterogeneity of the population and the bands of sender cells that form. A trap with a larger number of thin bands of each strain resulted in all receiver cells expressing significant amounts of CFP (Fig. 6b, and Fig. S13). However, in a trap with a few number of thick bands, the receiver cells reduced to background CFP expression with increased distance away from the sender-strain bands, as expected from our model. To estimate the limits on the signaling distance, we looked for an experimental image with adjacent “wide” bands of each strain in order to observe the isolated decay of CFP in the receiver strain as a function of distance from the sender-receiver strain boundary. We fit the decay of CFP in a narrow region near the middle of the trap to our model using an exponential decay with a single, spatial decay rate parameter *ξ*, and measured *ξ* = 20 μm (see example experimental receiver decay and exponential fit in Fig. 6d).

## Concluding remarks

Our results demonstrate that both the population size within and the geometry of a microfluidic trap can affect the spatiotemporal dynamics of microbial consortia. We have aimed in this study to characterize the ideal microfluidic trap environment for consortia by analyzing vital components such as strain ratio stability and quorum-sensing molecule signaling distance. Among the trap designs we tested, small three-walled traps allow for more spatially uniform signaling but suffer from strain ratio instability. By contrast, large open traps exhibit more robust strain ratio stability. Our quantification of effective signaling distance implies that strains must be reasonably well-mixed in order for inter-strain communication to function properly. We have shown both experimentally and through modeling that such spatial heterogeneity can be achieved by controlling the number of seed cells. Our experimental data also showed that increasing the number of seed cells in a large open trap can increase strain ratio instability, suggesting competing objectives for our experiments. However, the open traps with large number of seed cells (resulting in thin bands of cells ideal for communication) still have more stable populations than the hallway traps. Maintaining strain ratio stability and intercellular communication is vital for synthetic microbial consortia and our results suggest how to address this balance in the microfluidic environment.

**Figure 6:**
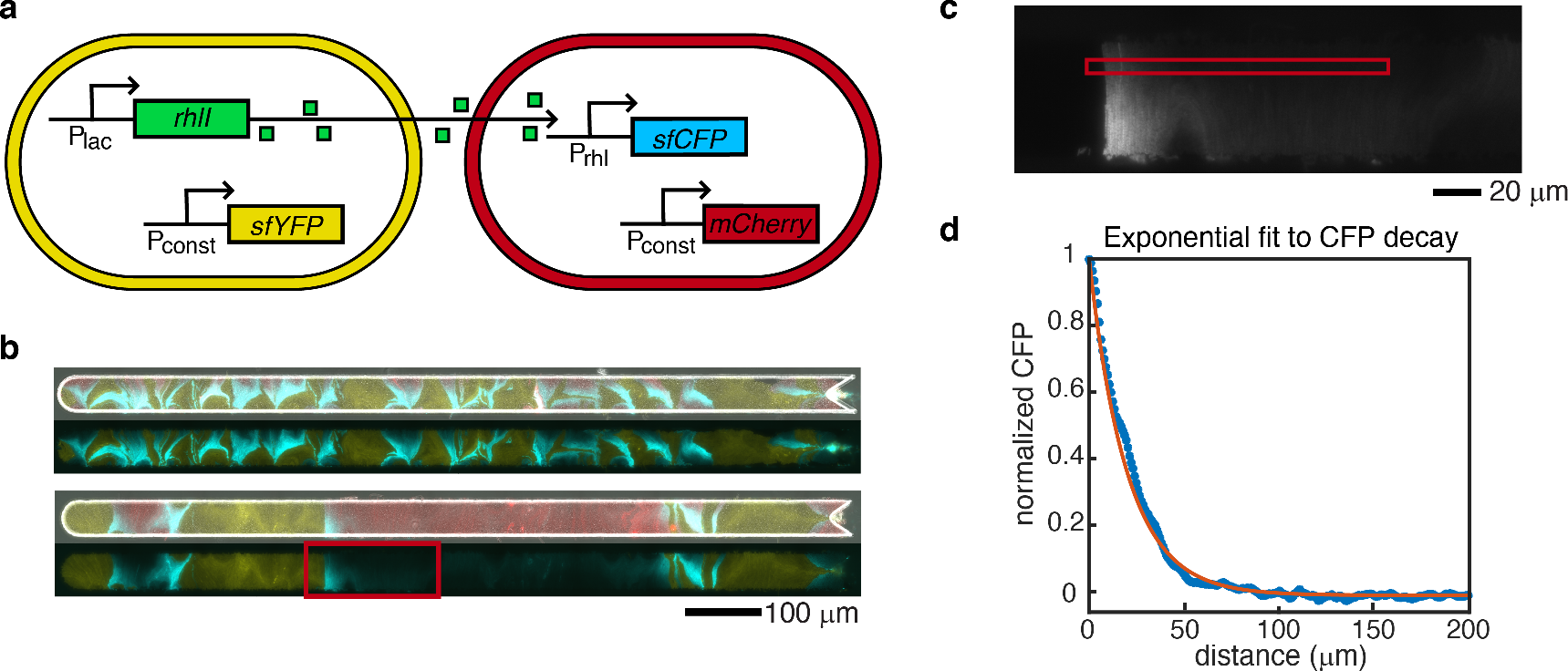
Sender-receiver consortium and signaling depth measurement. **(a)** Schematic of the sender and receiver strain gene circuits. Genes shown are on plasmids. **(b)** In the top pair of images the larger number of thinner, interspersed bands results in sufficient intermingling of strains so that all receiver cells respond to QS molecules. The smaller number of thicker single-strain bands in the lower image pair leaves some receiver cells at a large distance from the sender strain. As a result, CFP signal decreases to background levels as the distance from a sender band increases. Top images of each pair are merged phase contrast, and yellow, red, and cyan fluorescent images from the microscope. Bottom images of each pair are only yellow and cyan fluorescent images merged to more clearly visualize CFP decay. **(c)** We selected a subset of the cells in the experimental image in (b) to determine the decay of CFP signal with distance from the sender strain. We averaged the signal in the red square over columns, and in a series of ten frames forward from the shown image, to generate the experimental CFP signal. Transition boundary from left (sender cells) to right (receiver cells) was automatically detected and set to the x-axis (distance) origin. **(d)** After threshold detection for receiver cells, response data was normalized and fit to an exponential decay with spatial decay parameter *ξ*. Experimental data shown as blue stars, fit curve in orange. Resulting fit: *ξ* =20 μm.

## Methods

### Construction of non-communicating strains

The non-communicating strains contained plasmids built through PCR and restriction enzyme cloning. These plasmids contained ampicillin resistance and p15A origin. A constitutive P_Iq_ promoter^29^ and modified bicistronic design ribosome binding site^30^ drove the expression of *sfcfp* or *sfyfp* mutants of *sfgfp*.^21,22^ The fluorescent proteins were tagged with a mutagenized ssrA tag that ends with amino acids ASV.^31^ LAA, ASV, and untagged versions were constructed, sequence verified and then tested on a plate reader and microscope. ASV-tagged versions were selected because they had detectable expression on the plate reader and did not saturate the detector of the microscope camera. Downstream of the fluorescent protein gene is the iGEM registry B0014 terminator. The plasmids were individually transformed into BW25113 ΔaraC ΔlacI^3^ *E. coli* cells using chemically competent cells and a heat shock transformation protocol.

The transformations were plated onto LB agar plates with 100 μg mL^−1^ ampicillin and incubated 16-18 hours at 37 C. The next day a colony was picked from each to inoculate a 5 mL tube of LB broth with 100 μg mL^−1^ ampicillin and in a shaking incubator at 37 C for 16-18 hours. The next day the cultures were diluted 1/1000 each into 50ml fresh LB with 100 μg mL^−1^ ampicillin and grown about 2.5 hours until they reached an OD600 nm of 0.15 − 0.2. While the cells grew, the microfluidic device was warmed to 37 C then flushed with 1o/ Tween-20 to purge air. Then media and waste syringes were then prepped. A 20ml syringe with LB with 0.075% Tween-20 and 100 μg mL^−1^ ampicillin was attached to the media port of the microfluidic device. Two 10 mL syringes with sterile water were attached to the two waste ports of the microfluidic device. Once the cultures reached the proper OD, 15 mL of each culture was spun down at 2000 *xg* for 5 minutes at room temperature. The supernatant was removed, and the cells were resuspended in 10 mL total of LB with 100 μg mL^−1^ ampicillin. This media was pre-warmed to 37 C. The 10 mL of mixed cells was seeded into a 10 mL syringe and attached to the microfluidic device.

The heights of the four syringes determine the flow of media through the device. The waste syringes were at the lowest height, the cell syringe began at a level higher than the waste but lower than the media to allow for cell loading, then it was lowered to around the waste syringe height to allow the media to reach the cells. When loading cells into the open trap, cells flew into the “V” on the right-hand side of the device and the line was flicked to apply enough pressure for the cells to go into the trap. They were then stuck in the trap whose height 0.95 μm was slightly less than the diameter of the cells 1 μm. Once the desired number of cells was trapped, the cell syringe was lowered to allow media flow to the cells. The device was then moved to a 60X oil objective and imaged immediately every 6 minutes at phase contrast, YFP, and CFP filter settings. They were imaged immediately to capture the initial number of cells.

For the hallway trap, the loading height of the cell syringe was adjusted to allow the cells to flow into the traps on their own. It was left at this loading height for 30 minutes to an hour to trap as many cells as possible. This was the best way to ensure both strains entered the traps. Afterwards the cell syringe was again lowered to allow media flow to the cells. The cells were allowed to grow prior to imaging for 2-4 hours before moving to a higher (100X) oil objective and imaged immediately every 6 minutes at phase contrast, YFP, and CFP filter settings.

### Construction of sender and receiver strains

The sender strain used to test communication in the open trap had the same constitutive *sfyfp* plasmid used in the non-communicating strains above for identification. It also contained a plasmid encoding the *rhlI* (ATCC #47085) gene driven by a promoter under the control of LacI with the same bicistronic design ribosome binding site^30^ used previously. RhlI produces the C4-HSL QS molecule that can diffuse out the cell to nearby cells.^28^ The *rhlI* gene was tagged with the original LAA ssrA degradation tag^31^ and had the iGEM registry B0014 terminator. This plasmid contained a kanamycin resistance gene for selection, pMB1 origin, and ROP element that reduces the copy number.^32^ This plasmid was transformed into a BW25113 ΔaraC ΔlacI ΔsdiA strain with araC, cinR, and rhlR inserted at the attB site under constitutive promoter.^10^ There was no LacI in this strain, so rhlI was expressed constitutively. The receiver strain had a plasmid with the same backbone but contained an engineered promoter with an RhlR binding site driving the expression of *sfcfp*.^21,22^ The *sfcfp* gene was not tagged for degradation but has the same iGEM registry B0014 terminator. This plasmid was transformed into the same BV25113 ΔaraC ΔlacI ΔsdiA strain with araC, cinR and rhlR inserted at the attB site under constitutive promoter.^10^ The expression of *sfcfp* is dependent upon activation when RhlR binds C4-HSL^28^ produced by the sender or exogenously added to the media. For identification, the receiver contained a plasmid identical to the constitutive *sfyfp* plasmid used in the sender and non-communicating strains, but with mCherry^33^ in place of *sfyfp*. These strains were transformed, cultured, and tested in the microfluidic devices using the same protocol as the non-communicating strains described previously.

### Data analysis and microfluidic device construction

Individual fluorescent images obtained from the microscope Nikon Elements program were exported into tiff files for each time and channel – bit depth was conserved during export (12-bit). These images were analyzed using custom MATLAB code, and Ilastik machine learning software was used to identify cells for the strain ratio analyses. Images and videos were compiled in ImageJ. Wafer molds and the actual microfluidic devices were made using methods described in Ferry et al., 2011.^16^ The device with original hallway trap is the same device used in Chen et al., 2015,^10^ and the open trap device is based off the device from Hussain et al., 2014^34^ but with a rounded end and 2000 μm length. The varied length devices were constructed the same way as the original open and hallway traps, but with different length trap designs in the mask. For increased throughput, some open trap experiments were done using a parallel device we constructed which contains 4 of the original 2mm long open traps. This parallel device was designed off the biopixel device from Prindle et al., 2012.^19^ We reduced the number of parallel channels to four, removed their traps, and inserted our narrow open traps. We also used this schematic for the 4 reduced length open traps in which one of the four length traps was inserted into each of the four parallel channels resulting in a device with four open traps with lengths of 1mm, 0.5mm, 0.225mm, and 0.1mm.

## Acknowledgenent

This work was supported by the National Institutes of Health, through grant R01GM117138 (MRB, KJ, WO) and the joint NSF-National Institute of General Medical Sciences Mathematical Biology Program grant R01GM104974 (MRB, KJ, WO); NSF grants DMS-1413437 (WO), DMS-1662290 (MRB), and GRFP-1450681(RNA); and the Welch Foundation grant C-1729 (MRB).

The authors acknowledge the use of the Opuntia Cluster and support from the Center of Advanced Computing and Data Systems at the University of Houston.

## References

(1) Elowitz, M. B.; Leibier, S. A synthetic oscillatory network of transcriptional regulators. Nature 2000, 403, 335–338.

(2) Gardner, T. S.; Cantor, C. R.; Collins, J. J. Construction of a genetic toggle switch in Escherichia coli. Nature 2000, 403, 339–342.

(3) Stricker, J.; Cookson, S.; Bennett, M. R.; Mather, W. H.; Tsimring, L. S.; Hasty, J. A fast, robust and tunable synthetic gene oscillator. Nature 2008, 456, 516–519.

(4) Brenner, K.; You, L.; Arnold, F. H. Engineering microbial consortia: a new frontier in synthetic biology. Trends in Biotechnology 2008, 26, 483–489.

(5) You, L.; Cox, R. S.; Weiss, R.; Arnold, F. H. Programmed population control by cell-cell communication and regulated killing. Nature 2004, 428, 868–871.

(6) Nadell, C. D.; Drescher, K.; Foster, K. R. Spatial structure, cooperation and competition in biofilms. Nature Publishing Group 2016, 14, 589–600.

(7) Johns, N. I.; Blazejewski, T.; Gomes, A. L.; Wang, H. H. Principles for designing synthetic microbial communities. Current Opinion in Microbiology 2016, 31, 146–153.

(8) Jones, J. A.; Wang, X. Use of bacterial co-cultures for the effi cient production of chemicals. Current Opinion in Biotechnology 2018, 53, 33–38.

(9) Bittihn, P.; Din, M. O.; Tsimring, L. S.; Hasty, J. Rational engineering of synthetic microbial systems: from single cells to consortia. Current Opinion in Microbiology 2018, 45, 92–99.

(10) Chen, Y.; Kim, J. K.; Hirning, A. J.; Josić, K.; Bennett, M. R. Emergent genetic oscillations in a synthetic microbial consortium. Science 2015, 349, 986–989.

(11) Tsoi, R.; Wu, F.; Zhang, C.; Bewick, S.; Karig, D.; You, L. Metabolic division of labor in microbial systems. Proceedings of the National Academy of Sciences 2018, 115, 201716888.

(12) Shong, J.; Jimenez Diaz, M. R.; Collins, C. H. Towards synthetic microbial consortia for bioprocessing. Current Opinion in Biotechnology 2012, 23, 798–802.

(13) Potvin-Trottier, L.; Luro, S.; Paulsson, J. Microfluidics and single-cell microscopy to study stochastic processes in bacteria. Current Opinion in Microbiology 2018, 43, 186–192.

(14) Kou, S.; Cheng, D.; Sun, F.; Hsing, I.-M. Microfluidics and microbial engineering. Lab Chip 2016, 16, 432–446.

(15) Bennett, M. R.; Hasty, J. Microfluidic devices for measuring gene network dynamics in single cells. Nature Reviews Genetics 2009, 10, 628.

(16) Ferry, M.; Razinkov, I.; Hasty, J. Synthetic Biology, Part A, 1st ed.; Elsevier Inc., 2011; Vol. 497; pp 295–372.

(17) Razinkov, I. A.; Baumgartner, B. L.; Bennett, M. R.; Tsimring, L. S.; Hasty, J. Measuring competitive fitness in dynamic environments. Journal of Physical Chemistry B 2013, 117, 13175–13181.

(18) Danino, T.; Mondragón-Palomino, O.; Tsimring, L.; Hasty, J. A synchronized quorum of genetic clocks. Nature 2010, 463, 326–330.

(19) Prindle, A.; Samayoa, P.; Razinkov, I.; Danino, T.; Tsimring, L. S.; Hasty, J. A sensing array of radically coupled genetic ‘biopixels’. Nature 2012, 481, 39–44.

(20) Bacchus, W.; Fussenegger, M. Engineering of synthetic intercellular communication systems. Metabolic Engineering 2013, 16, 33–41.

(21) Kremers, G. J.; Goedhart, J.; Van Munster, E. B.; Gadella, T. W. Cyan and yellow super fluorescent proteins with improved brightness, protein folding, and FRET förster radius. Biochemistry 2006, 45, 6570–6580.

(22) Pèdelacq, J. D.; Cabantous, S.; Tran, T.; Terwilliger, T. C.; Waldo, G. S. Engineering and characterization of a superfolder green fluorescent protein. Nature Biotechnology 2006, 24, 79–88.

(23) Boyer, D.; Mather, W.; Mondragón-Palomino, O.; Orozco-Fuentes, S.; Danino, T.; Hasty, J.; Tsimring, L. S. Buckling instability in ordered bacterial colonies. Physical Biology 2011, 8, 026008.

(24) Winkle, J. J.; Igoshin, O. A.; Bennett, M. R.; Josić, K.; Ott, W. Modeling mechanical interactions in growing populations of rod-shaped bacteria. Physical Biology 2017, 14.

(25) Dell’Arciprete, D.; Blow, M. L.; Brown, A. T.; Farrell, F. D. C.; Lintuvuori, J. S.; McVey, A. F.; Marenduzzo, D.; Poon, W. C. K. A growing bacterial colony in two dimensions as an active nematic. Nature Communications 2018, 9, 4190.

(26) Volfson, D.; Cookson, S.; Hasty, J.; Tsimring, L. S. Biomechanical ordering of dense cell populations. Proceedings of the National Academy of Sciences of the United States of America 2008, 105, 15346–15351.

(27) Karamched, B. R.; Ott, W.; Timofeyev, I.; Alnahhas, R. N.; Bennett, M. R.; Josić, K. Moran model of spatial alignment in microbial colonies. Physica D: Nonlinear Phenomena 2019, https://doi.org/10.1016/j.physd.2019.02.001.

(28) Pesci, E. C.; Pearson, J. P.; Seed, P. C.; Pesci, E. C.; Pearson, J. P.; Seed, P. C.; Iglewski, B. H. Regulation of las and rhl quorum sensing in Pseudomonas aeruginosa Regulation of las and rhl Quorum Sensing in Pseudomonas aeruginosa. Strain 1997, 179, 3127–3132.

(29) Calos, M. P. DNA sequence for a low-level promoter of the lac repressor gene and an ‘up’ promoter mutation. Nature 1978, 274, 762–765.

(30) Mutalik, V. K.; Guimaraes, J. C.; Cambray, G.; Lam, C.; Christoffersen, M. J.; Mai, Q. A.; Tran, A. B.; Paull, M.; Keasling, J. D.; Arkin, A. P.; Endy, D. Precise and reliable gene expression via standard transcription and translation initiation elements. Nature Methods 2013, 10, 354–360.

(31) Andersen, J. B.; Sternberg, C.; Poulsen, L. K.; Bjørn, S. P.; Givskov, M.; Molin, S. New Unstable Variants of Green Fluorescent Protein for Studies of Transient Gene Expression in Bacteria New Unstable Variants of Green Fluorescent Protein for Studies of Transient Gene Expression in Bacteria. Applied and environmental microbiology (1998) 1998, 64, 2240–2246.

(32) Lacatena, R. M.; Banner, D. W.; Castagnoli, L.; Cesareni, G. Control of initiation of pMB1 replication: Purified rop protein and RNA I affect primer formation in vitro. Cell 1984, 37, 1009–1014.

(33) Shaner, N. C.; Campbell, R. E.; Steinbach, P. A.; Giepmans, B. N.; Palmer, A. E.; Tsien, R. Y. Improved monomeric red, orange and yellow fluorescent proteins derived from Discosoma sp. red fluorescent protein. Nature Biotechnology 2004, 22, 1567–1572.

(34) Hussain, F.; Gupta, C.; Hirning, A. J.; Ott, W.; Matthews, K. S.; Josic, K.; Bennett, M. R. Engineered temperature compensation in a synthetic genetic clock. Proceedings of the National Academy of Sciences 2014, 111, 972–977.

