## Supplemental Information for "Spatiotemporal dynamics of synthetic microbial consortia in microfluidic devices"

### Supporting Information Available

#### Coin-flipping Model

To characterize the expected number of strain bands in the "open" microfluidic device, we used a sequential "coin-flipping" model for the initial seed strain and placement of cells, where one of two strains was chosen at random and placed with equal probability along the centerline of the trap (this modeled the "loading" of cells in the trap). We assumed a one-dimensional reduction of the initial seed positions due to the large aspect ratio ($\approx 20:1$) of the open device. We further assumed that initial seed cells grew symmetrically until making contact with another seed cell colony; at which time, if the two colonies were of different strain types, the merged colony would form a strainA-strainB interface. The colonies in this way would continue to expand (possibly forming other interfaces with other single or merged colonies) until the trap filled. We also assumed that a full trap resulted in vertical, single cell-width columns, where the identity of each column was of strain type 'A' or 'B' (in the experimental data these were sender or receiver cells) and that each column was derived directly from an initial seed cell colony. Cells were thus modeled as being initialized in the middle of the trap and then growing uniformly until the trap consisted completely of side-by-side columns.

We are interested in the expected number of *bands* of cells vs. the number of cells seeded, where a band is a contiguous group of columns, all of the same strain type. The number of bands is determined by the number of strainA-strainB interfaces that form during the colonies' expansion phase after initial cell seeding. Let $n$ be the number of seed cells (seeded with equal probability 0.5 of each strain type) and let $b$ represent the number of resulting bands. The expectation and variance of the number of bands is computed as a function of the number of seed cells $n$ as:

$$\mathbb{E}[b] = \frac{n+1}{2} \qquad \mathbf{var}(b) = \frac{n-1}{4}. \tag{1}$$

We used a range of $\pm 3$ standard deviations in Figure 5 that is based on the above calculation for each number of initial seed cells.

#### Diffusion Model

Let $x \in \mathbb{R}^+ \equiv [0, \infty)$. Partiton $\mathbb{R}^+$ into two subsets $I_1, I_2$, where $I_1 = [0, L_s]$ and $I_2 = (L_s, \infty)$. $I_1$ will correspond to the region where we have a densely packed sender strain. Hence, $L_s$ represents the thickness of the stripe formed by these sender cells. $I_2$ will be the region where receiver cells, for example, may exist. We are interested in understanding the density of a chemical signal $U$ produced by the sender strain uniformly in $I_1$ at a point that is a distance $\delta$ away from the right endpoint of $I_1$ (i.e. a distance $\delta$ from the stripe of sender cells) as a function of the thickness of the stripe, $L_s$. Let $u_1(x,t)$ denote the density of chemical $U$ at a point $x$ at time $t$ in $I_1$. Let $u_2$ describe the corresponding density in $I_2$. The following are the dynamics for $u_1, u_2$:

$$\frac{\partial u_1}{\partial t} = \alpha + D\frac{\partial^2 u_1}{\partial x^2} - \gamma u_1, \quad x \in I_1 \tag{2}$$

$$\frac{\partial u_2}{\partial t} = D\frac{\partial^2 u_2}{\partial x^2} - \gamma u_2, \quad x \in I_2 \tag{3}$$

The signal diffuses and degrades at a rate $\gamma$ in all of $\mathbb{R}^+$ but in $I_1$ the signal is produced at some rate $\alpha$. This is what is captured in these equations. We note that $\gamma$ represents a degradation in a loose sense. Most likely this rate will represent a rate of absorption or consumption of the signal molecule. The boundary conditions are:

$$\left.\frac{\partial u_1}{\partial x}\right|_{x=0} = 0$$

$$\lim_{x \to \infty} u_2 = 0$$

$$u_1(L_s, t) = u_2(L_s, t)$$

$$\left.\frac{\partial u_1}{\partial x} = \frac{\partial u_2}{\partial x}\right|_{x=L_s}$$

The first boundary condition says none of the chemical exits from the boundary at $x = 0$. The second boundary condition keeps the solutions physical by preventing blowup. The last two boundary conditions impose continuity in the density and first derivative of the density at the partitioning point $x = L_s$. Examining this system at steady state and imposing the first two boundary conditions, we obtain

$$u_1(x) = 2A\cosh\left(x\sqrt{\frac{\gamma}{D}}\right) + \frac{\alpha}{\gamma} \tag{4}$$

$$u_2(x) = Be^{-x\sqrt{\frac{\gamma}{D}}} \tag{5}$$

The last step is to impose continuity at $x = L_s$. In doing this, we obtain the linear system

$$\mathbf{A}\mathbf{v} = \mathbf{b},$$

where

$$\mathbf{A} = \begin{bmatrix} 2\cosh\left(L_s\sqrt{\frac{\gamma}{D}}\right) & e^{-L_s\sqrt{\frac{\gamma}{D}}} \\ 2\sinh\left(L_s\sqrt{\frac{\gamma}{D}}\right) & e^{-L_s\sqrt{\frac{\gamma}{D}}} \end{bmatrix}, \mathbf{v} = \begin{bmatrix} A \\ B \end{bmatrix}, \mathbf{b} = \begin{bmatrix} -\frac{\alpha}{\gamma} \\ 0 \end{bmatrix}$$

We now plot $u_2(L_s + \delta)$ as a function of $L_s$. We can also plot the density profile of the chemical signal $u(x)$. It is also straightforward to generalize the above framework to the case where there are receiver cells on either side of the region where sender cells are densely packed. In this case, we let our domain be all of $\mathbb{R}$ and partition it into $I_1, I_2, I_3$, where $I_1 = (-\infty, -L_s/2)$, $I_2 = [-L_s/2, L_s/2]$, and $I_3 = (L_s/2, \infty)$. We let $u_i(x,t)$ be the density of chemical $U$ in the set $I_i$, for $i = 1, 2, 3$. We impose zero conditions at $x = \pm\infty$ and continuity conditions analogous to the above case at $x = \pm\frac{L_s}{2}$.

$$\frac{\partial u_1}{\partial t} = D\frac{\partial^2 u_1}{\partial x^2} - \gamma u_1, \quad x \in I_1 \tag{6}$$

$$\frac{\partial u_2}{\partial t} = \alpha + D\frac{\partial^2 u_2}{\partial x^2} - \gamma u_2, \quad x \in I_2 \tag{7}$$

$$\frac{\partial u_3}{\partial t} = D\frac{\partial^2 u_3}{\partial x^2} - \gamma u_3, \quad x \in I_3 \tag{8}$$

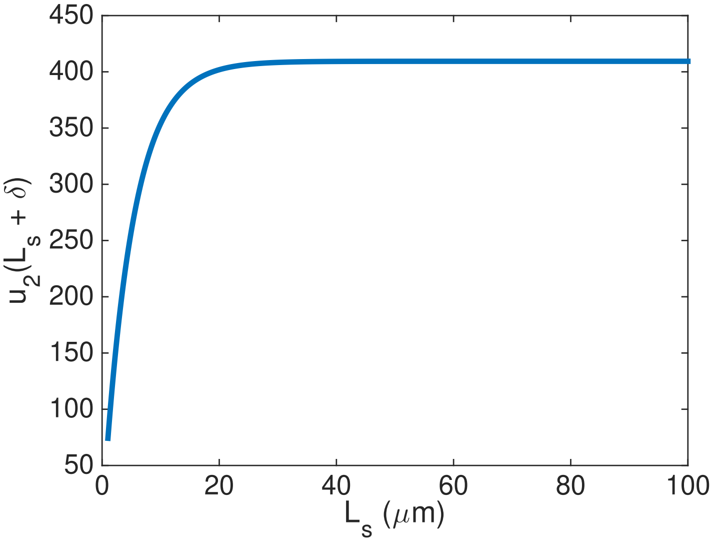


Figure S1: Plot of $u_2(L_s + \delta)$ as a function of $L_s$. Parameter values are $D = 1$, $\gamma = 0.01$, $\delta = 2$, and $\alpha = 10$.

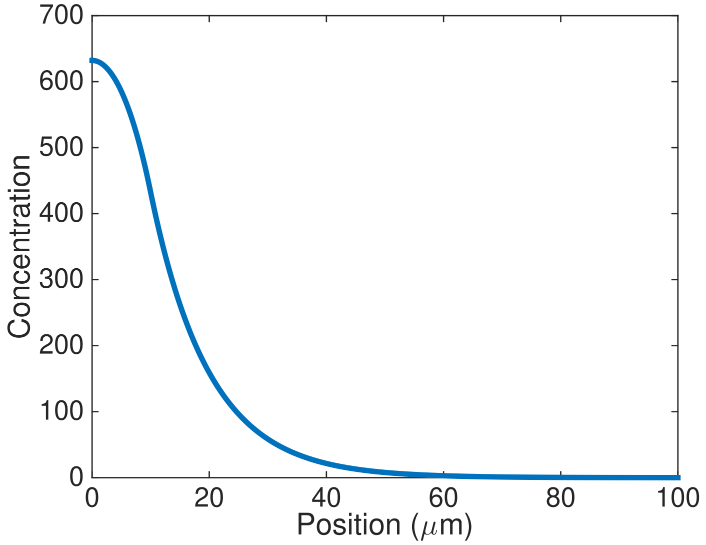


Figure S2: Plot of $u(x)$. Parameter values are $D = 1$, $\gamma = 0.01$, $L_s = 10$, and $\alpha = 10$.

We study this system at steady state. Imposing continuity boundary conditions yields a $4 \times 4$ linear system analogous to the $2 \times 2$ system solved in the one-sided case. The key

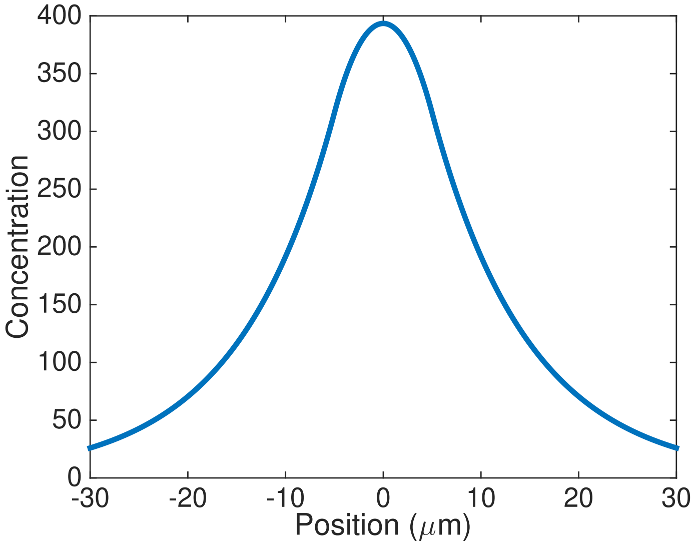


Figure S3: Plots of $u(x)$ in the symmetric case. Parameter values are $D = 1$, $\gamma = 0.01$, $L_s = 10$, and $\alpha = 10$.

parameter in all these results is the spatial correlation length, $\xi \equiv \sqrt{D/\gamma}$, along which the decay of the signal molecule produced by the sender strain occurs. Figure S1 shows that for $0 < L_s < \xi$, the density of signal felt at an area outside the stripe of sender cells increases as the thickness of the stripe increases. On the other hand, for $L_s > \xi$, there is not much change in the density of signal felt outside the stripe. Hence there is an optimal value for the thickness of the sender stripe, given by $L_s = \xi$, where a sender stripe can maximize its range of influence at a minimal metabolic load. Figures S2 and S3 show that the decay of the spatial profile of the signal molecule occurs with a characteristic length scale given by $\xi$. This simple framework provides us with a means to look at several things. Namely, it gives an estimate to how far a receiver cell can be from a sender cell and respond to the signal molecule.

#### Supplementary Figures

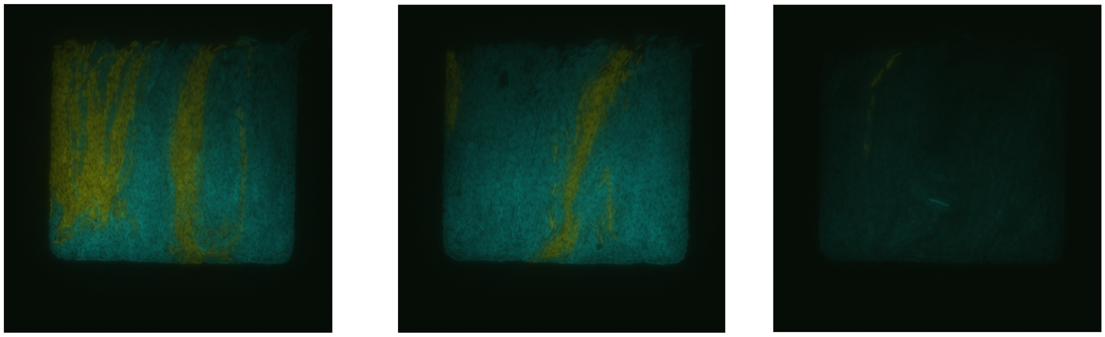

Figure S4: **Signaling in the small hallway trap.** Images of sender cells (yellow) cultured with receiver cells (cyan) over time in a hallway trap. Receiver cells fluoresce cyan in the presence of C4 HSL produced by sender cells. These images show that all receiver cells fluoresce cyan whenever there are sender cells present - up until the loss of sender cells in the trap in the rightmost image.

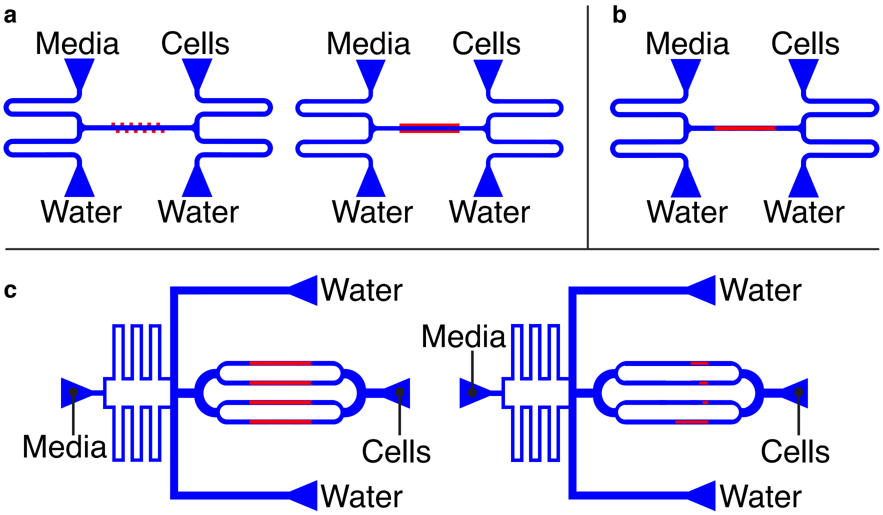


Figure S5: **Schematic of the entire microfluidic devices.** Blue regions are the 10 micron tall flow channel and the red regions are the cell-trapping regions. Blue triangle are ports to which media, cells, and waste reservoirs are connected. **(a)** The original hallway (left) and extended hallway (right) trap devices. Traps (red) are 1.5 microns tall. **(b)** The original(2mm) open trap device. Traps (red) are 0.95 microns tall. **(c)** The parallel device with four 2mm long open traps or with four different length traps: 1mm, 0.5mm, 0.225mm, and 0.1mm. Traps (red) are 0.95 microns tall.

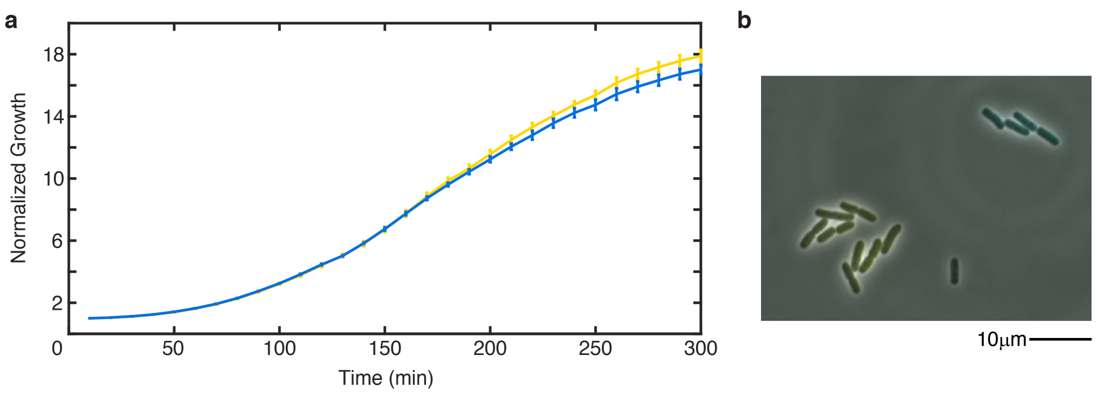


Figure S6: **Non-communicating strains have equal growth rates and sizes. (a)** Growth experiments in a 96-well plate over time show no significant difference in the growth rate of the two non-communicating strains. **(b)** Microscope images show similar size and shape of non-communicating strain in the fluidic devices.

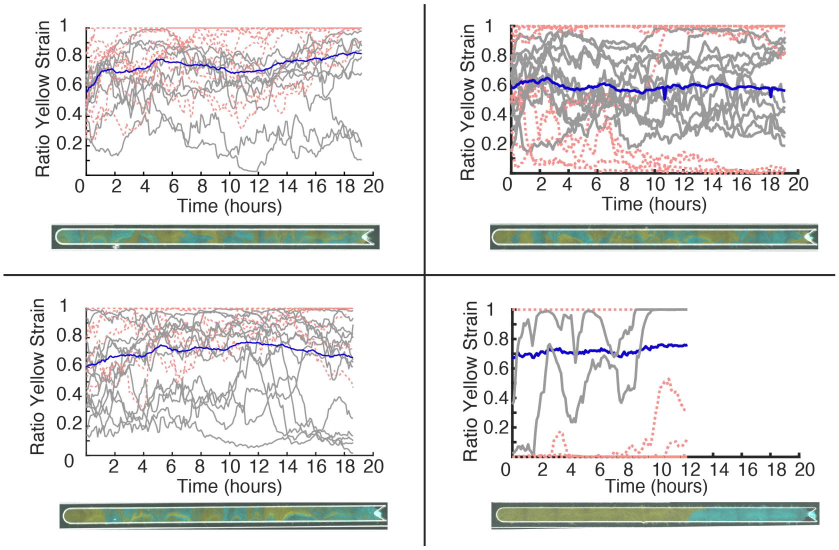


Figure S7: **Segmented open trap examples.** Data from four additional open traps when segmented. Image of each trap once filled with cells is shown below each graph. Blue lines are the entire trap yellow strain ratio; gray solid and red dashed lines are data from each hallway-length segment. Red dashed lines are segments that at some point lose one strain. All data looks noisy with hallway-like strain instability except the bottom right which shows a trap with only has about 2 bands of cell strains.

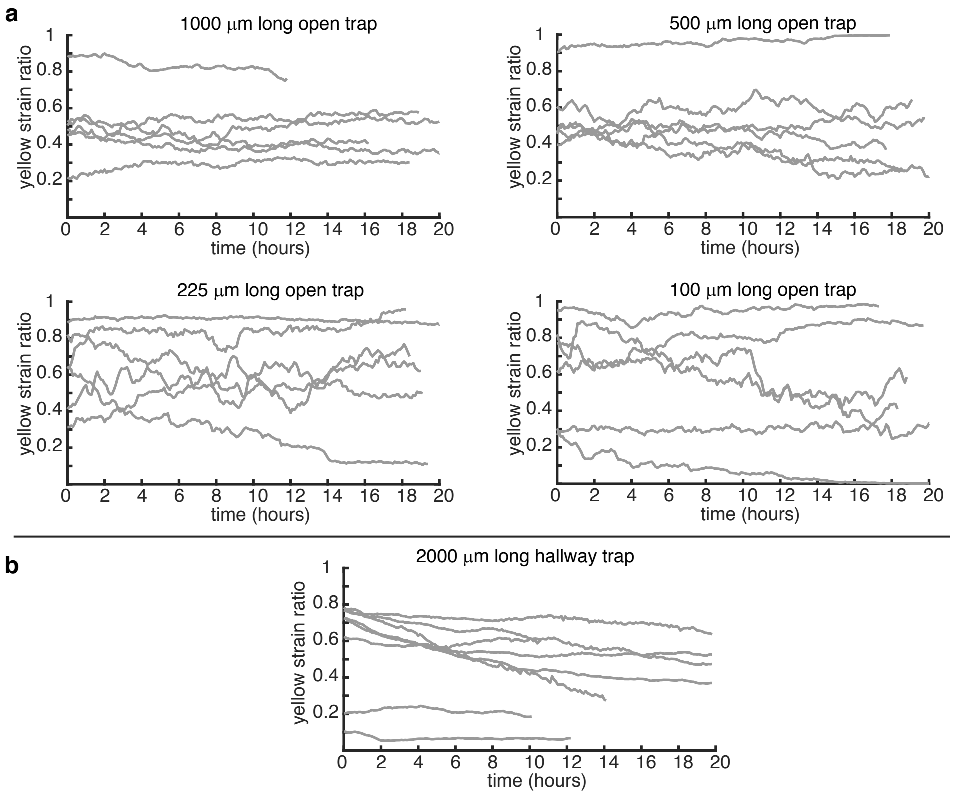


Figure S8: **Time Series from varied length traps.** Data of strain ratio over time from each varied length trap experiment averaged in Figure4. **(a)** As the open trap gets shorter, the strain ratios become more variable over time. **(b)** The longer (2000 µm) hallway trap shows more stability than the original length (100 µm) hallway data, but shows significant drift.

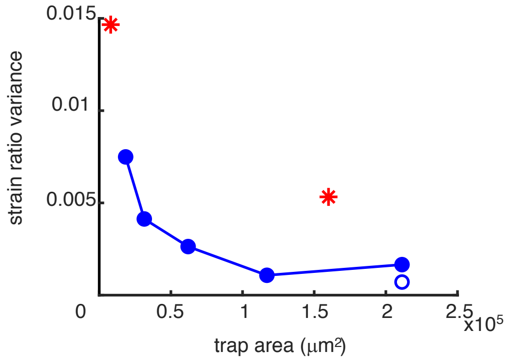


Figure S9: **Average strain ratio time series variance.** The variance of strain ratio over time is shown to compare with Figure 4 in the main text, which plots the range of strain ratios. The two plots are qualitatively the same and show the trend of increased strain ratio stability with increasing size of the cell-trapping area. Closed blue circles: average strain ratio variance of the open-walled devices of each trap size. The open circle for the longest open-walled device is lowered (open blue circle) when including only experiments with the 6 smallest number of resulting stripe bands (see main text and Figures S10 and S11) Red stars: same data but for the "hallway" device of the two experimental sizes, which show significant deviation from the extended data.

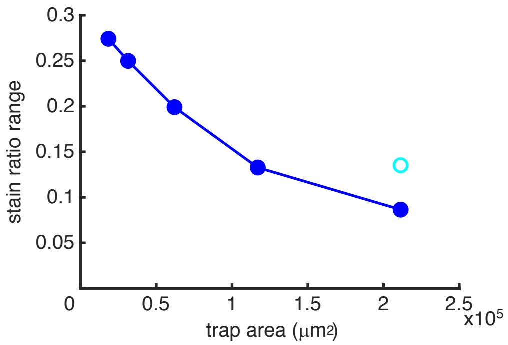


Figure S10: **Average strain ratio range vs. trap area for reduced data in the longest trap size.** We reduced the data set in the longest open-walled trap to compare with Figure 4 in the main text. The reduced data reflect a similar number of striping bands for the experiment. Filled blue circles: mean strain ratio range of the open-walled devices of each trap size (x-axis: cell-occupied trap area) with the last data point including the reduced data set only. Open cyan circle: Mean range while including all data for the longest trap size (same as in Figure 4). Increasing numbers of seed cells correlated with increasing number of measured striping bands in the experiment, which, in turn, correlates with measured strain ratio variability. See Figure S11).

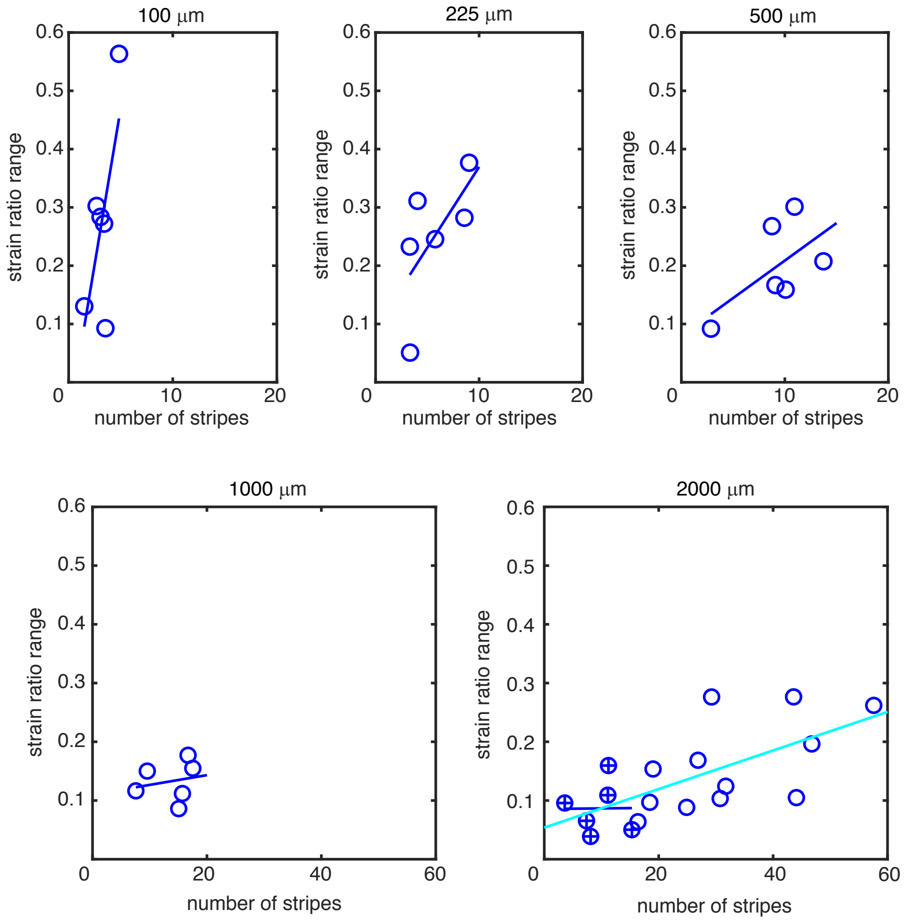


Figure S11: **Experimental strain ratio range vs. measured number of striping bands for each trap size.** Data for each length of open-walled "extended" trap showing measured strain ratio range vs. measured number of striping bands for each experiment in each device. Solid lines are linear least-squares fit of the data. In the 2000 μm length device, open blue circles show the entire data set, circles with filled + show the reduced data set, chosen to comprise the 6 data points with the smallest number of striping bands (thus, comparable to the data from other trap lengths). Blue line: least-squares linear fit to the reduced data. Cyan line: least-squares linear fit for the entire data set.

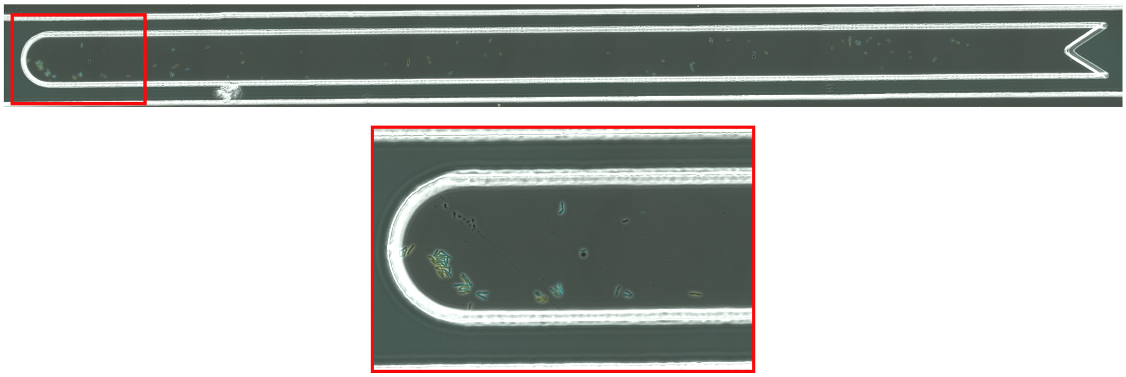

Figure S12: **Seeded cells clumped together.** This image is of the cells seeded into the last open trap from Figure 5 that does not fit into the model. As can be seen in the zoom in of the far left hand side of the trap, many cells have clumped together during the attempt to load a high number of cells into the trap. About 40 of the 123 cells seeded into the trap are in this small segment of the trap. Cells that are already clumped together when seeded will effectively act like one cell or colony and form one total band rather then each from their own band of cells when growing and filling the trap. This can explain why we observed fewer bands in this experiment than expected.

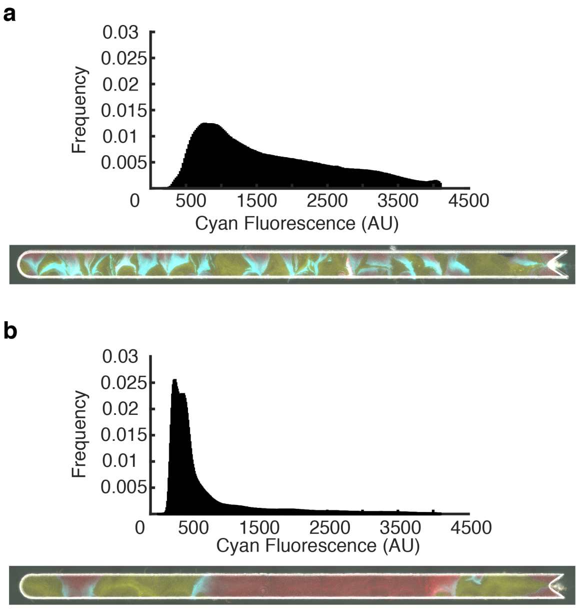


Figure S13: **Signalling in open trap.** **(a)** Distribution of CFP intensity of receiver cells in a well mixed open trap. All cells have fluorescence levels above background. **(b)** Distribution of CFP intensity of receiver cells in a less mixed open trap. Receiver cells further away from sender cells do not express significant levels of CFP.
